# Untapped viral diversity in global soil metagenomes

**DOI:** 10.1101/583997

**Authors:** Emily B. Graham, David Paez-Espino, Colin Brislawn, Kirsten S. Hofmockel, Ruonan Wu, Nikos C. Kyrpides, Janet K. Jansson, Jason E. McDermott

## Abstract

Viruses outnumber every other biological entity on Earth, and soil viruses are particularly diverse compared to other habitats. However, we have limited understanding of soil viruses because of the tremendous variation in soil ecosystems and because of the lack of appropriate screening tools. Here, we determined the global distribution of more than 24,000 soil viral sequences and their potential hosts, including >1,600 sequences associated with giant viruses. The viral sequences, derived from 668 terrestrial metagenomes, greatly extend existing knowledge of soil viral diversity and viral biogeographical distribution. We screened these sequences to identify a suite of cosmopolitan auxiliary metabolic genes (AMGs) encoding enzymes involved in soil organic carbon decomposition across soil biomes. Additionally, we provide evidence for viral facilitation of multi-domain linkages in soils by locating a fungal chitosanase in bacteriophages, generating a new paradigm of how viruses can serve as exchange vectors of carbon metabolism across domains of life.

---

Viruses outnumber every other biological entity on Earth by a wide margin^1^. Estimates suggest that 10^31^ virus particles exist globally, equivalent to the biomass of 75 million blue whales (200 million tons)^2^. A global meta-analysis of viral distribution revealed that the vast majority of viruses are clearly habitat-specific^3^. The soil virome in particular is poorly characterized in terms of its size and composition, but limited evidence shows that soil viruses are more abundant and diverse than viruses from other ecosystems^4,5,6^. This high viral diversity may be a result of the heterogeneous physical matrix of soil where spatial structuring generates a plethora of environmental niches^7,8^. Although viruses are recognized as key players in C and nutrient cycles in aquatic ecosystems^9^, we know comparatively little about the roles of viruses in soils. Major limitations to studies of soil viral ecology include difficulties in isolating soil viruses, the enormous range of soil ecosystems with distinct properties that prevent generalization between sites, and, until lately, the lack of appropriate molecular screening tools.

Shotgun metagenomics is a useful approach for analyzing soil viromes because most viruses lack a universal marker gene to target with primer-based methods^10^ and because many viruses in soil are double-stranded DNA phages that can be sequenced^11^. Several recent studies have used shotgun metagenomics in soil to characterize the soil virome in a limited range of soil types and biomes^12-16^. In the most comprehensive examination of soil viruses to date, exploration of 197 metagenomes from thawed permafrost enabled discovery of 3,112 soil viral sequences^12^. Emerson et al.^12^ predicted 14 viral glycoside hydrolase (GH) enzymes in these sequences with projected functions for breaking glycosidic linkages in pectin, hemicellulose, starch, and cellulose molecules; one of these was confirmed to express an endomannanase enzyme^12^. However, viral GHs can also be involved in general lysogenic functions (rev. in Davies et al.^17^), and the exact role of viral GHs in soils in unclear. Additionally, new nucleocytoplasmic large DNA viruses (NCDLV), colloquially known as giant viruses, were recently identified using a ‘mini-metagenomics’ approach in soil collected from the Harvard forest long-term experimental research site (LTER)^18^. To our knowledge, no study has investigated soil viruses at more than a few locations (Extended Data Fig. 1), and we lack a comprehensive assessment of virus diversity and function in soils, including the discovery of new viruses and their distribution, host associations, and possible exchange of genetic information across host domains.

Here, we greatly extend existing knowledge to identify >24,000 soil viral sequences from a wide range of globally distributed soil metagenomes. By deep analysis of the viral sequence data, we determine viruses that are prevalent across soil biomes, as well as their associations with soil microorganisms and patterns through space. In addition, we estimate the global distribution of soil viruses and identify thousands of new virally encoded auxiliary metabolic genes (AMGs) that could play key functional roles in soil ecosystems.

## Viral Abundance and Diversity

To assess the geographic distribution, diversity, and functional potential of viruses in soil microbiomes, we gathered metagenomic sequences from 668 soil samples spanning 75 locations on three continents (Supplementary Tables 1-2, Fig. 1A). We manually assigned biomes to each study using metadata deposited into Integrated Microbial Genomes with Microbiomes (IMG/M)^19^. We applied our previously described pipeline for identification of viral contiguous sequence regions (contigs)^3,20^ to the samples and identified 24,335 viral sequences (clustering into 17,229 unique viral operational taxonomic units, vOTUs) greater than 5 kilobases (kb), of which 20,700 were predicted to be bacteriophages, 96 were putative Eukaryotic viruses, and 3,300 were unknown (Supplementary Table 3). These viral sequences encoded a total of 4,306 distinct protein families (pfams) when searched against the Pfam database^21^ using HMMER 3.0^22^. Because many viral functions lack annotation in the Pfam database, we also conducted *de novo* sequence clustering to identify novel protein families which yielded 105,730 distinct clusters. Of these clusters, only 10,441 contained at least one function annotated in the Pfam database, indicating a large proportion of the functional capacity of soil viruses is completely novel. Additionally, we identified 1,676 sequences attributed to giant viruses and 538 sequences attributed to virophages of all sizes. Our study thus represents an enormous expansion of knowledge of the global distribution of soil viruses with a 25-fold increase in sample locations and a doubling of the number of soil viral sequences obtained (Extended Data Fig. 1).

**Figure 1.**
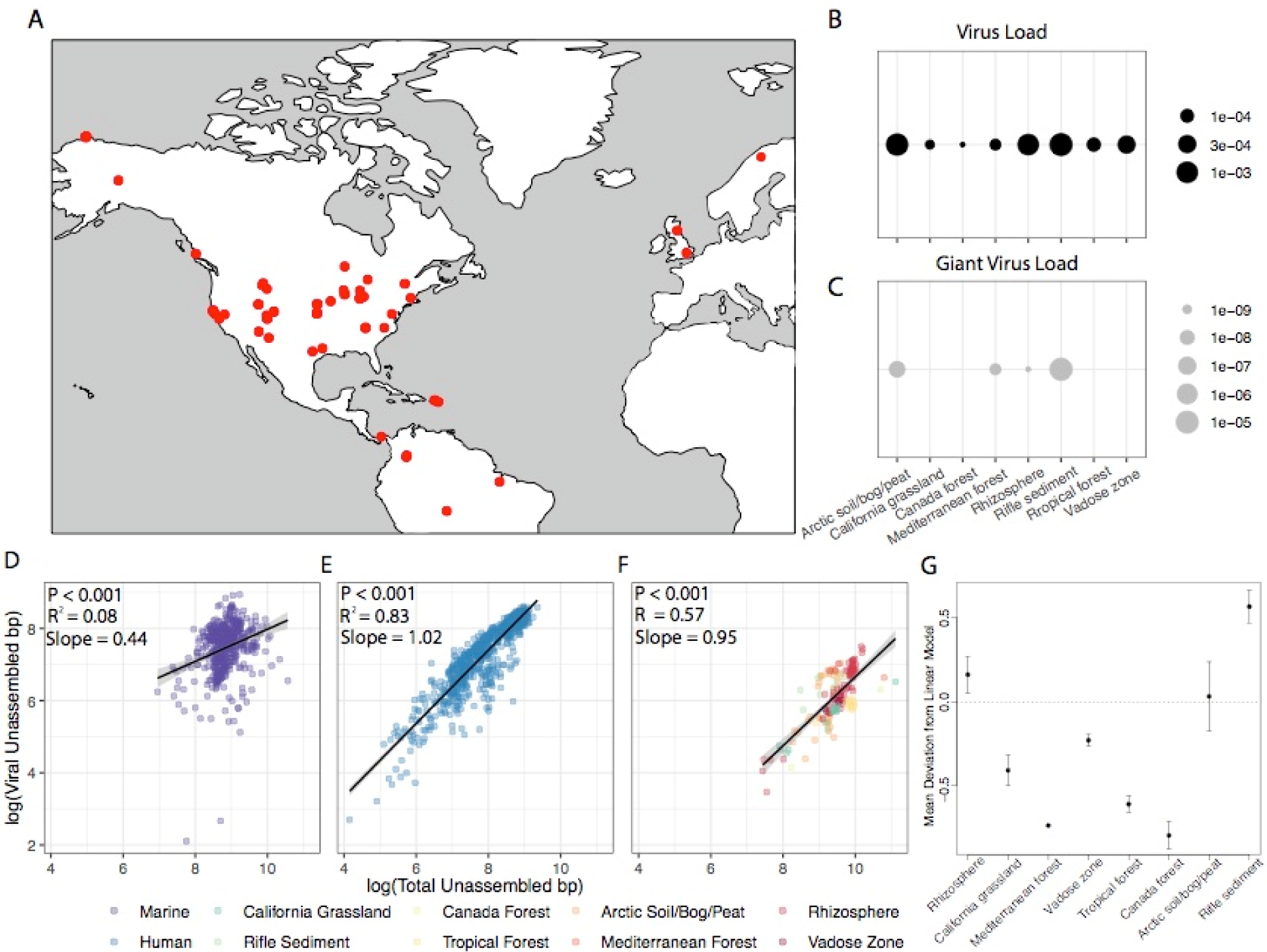
Soil metagenome locations and virome biogeography. Metagenomic sequences were obtained from 668 soil samples spanning 75 global locations. (A) Locations are denoted in red. Normalized abundances of sequences attributed to viruses and giant viruses are presented in (B) and (C) respectively. Normalized abundances were calculated at the ratio of viral sequence bp to total metagenomic bp, which showed log-linear relationships for (D) marine, (E) human, and (F) soil ecosystems. Colors in (F) represent the source biome of each sample. Deviations of each soil type from the expected viral load are presented in (G).

We assessed the relationship between metagenome size and number of viruses by comparing the number of base pairs from raw sequence reads that were attributed to viruses, versus all base pairs (bp) in the corresponding metagenome. This relationship was significant (P < 0.001, R^2^ = 0.58; Fig. 1F) with approximately 1 of every 5000 bp (0.2%) mapping to a viral sequence. Given an estimated average soil bacterial genome size of 5Mb, average soil viral genome size of 5000 bp^3^, and 10^9^ bacterial cells per gram dry soil, this equates to 2.37 × 10^8^ viruses per gram dry soil, in line with previous estimates of 10^7^ to 10^10^ viruses per gram dry weight soil^4,5,6^ and 10-100x fold lower than estimates in marine and human systems respectively (Fig. 1D-E). Fitting a quadratic model to the relationship between bacterial and viral bp did not improve fit, indicating that deeper sequencing will linearly increase the number of viruses discovered in soils and that soil viral diversity has been significantly under sampled by existing metagenomic sequencing efforts.

Subsequently, we explored the soil virome to examine biogeographical patterns in viral distributions. We sorted the metagenomes into 18 different biomes based on their ecosystem type in the GOLD database^23^ and Supplementary Table 2. We used the linear regression line between soil viral bp and microbial bp to identify biomes that had higher or lower abundances of viruses than expected (Fig. 1B, G). Rhizosphere soils and some terrestrial sediments had significantly higher viral loads than expected, whereas grassland and forest soils had lower viral concentrations (Fig. 1B, G, p ≤ 0.0001). Most soil viruses were highly biome-specific, but we identified 30 sequences that were cosmopolitan across grassland, forest, arctic, and rhizosphere biomes (Extended Data Table 1). Twelve of these sequences clustered into a single vOTU, and 14 sequences contained a single protein family (pfam00877) that is involved in phage lytic functions, providing an avenue for future investigation into viral traits that may be of particular importance in the soil virome.

Eukaryotic viruses with large genomes typically spanning several megabases have been identified in aquatic systems^24,25^ and, more recently, in terrestrial ecosystems, including soils^18,26,27^. Schulz et al.^18^ uncovered 16 novel giant virus genomes from metagenomes of forest soils and indicated that their discoveries constituted only a small fraction of the heretofore unknown giant virus diversity of soils. Here, we identified 1,676 putative giant virus sequences containing hallmark protein families of major capsids in NCLDV viruses (Supplementary Table 4). By filtering putative sequences to those with a contig length of greater than 5000 bp, we generated a list of 42 sequences that were assigned as giant viruses (longest assembled sequence of 259,840 bp). These 42 sequences were present in 19 samples – one rhizosphere, three arctic soils, one Mediterranean forest soil, three unassigned biomes, and eleven aquifer sediments from Rifle, Colorado. Additionally, the normalized abundance of giant viruses was 1-3 times greater in magnitude in Rifle samples than in any other sample set (Fig 1C).

Finally, 538 metagenomic viral contigs were assigned to virophages, DNA viral genomes that replicate along with giant viruses and co-infect eukaryotic cells, from which 26 were larger than 5000 bp; three of them were predicted to be complete (Supplementary Table 5).

## Eco-Evolutionary Patterns

Implementing ecological frameworks to understand biogeographic patterns has bolstered the growth of microbial ecology over the last few decades^28^. Such frameworks rely on commonly observed trends – such as latitudinal decreases in biomass and diversity^29^ and inverse correlations between community composition and geographic distance^30^ – to derive expectations for new environments and have aided in disentangling mechanisms driving biodiversity across the globe^31^. Given this history, we sought to uncover biogeographical patterns in the soil virome that could link soil viral diversity to established ecological frameworks.

Because of the wide range of geographic distances between our samples, we specifically focused on distance-decay relationships whereby community dissimilarity tends to increase with increasing distance^30^, and we hypothesized that phylogenetic distance between viruses would follow the same trend. Accordingly, we constructed *de novo* viral protein clusters through all-vs-all pairwise alignment of all open reading frames (Fig. 2A) and generated a phylogenetic tree for each viral protein clusters with at least 15 members (10,544). Over ten percent (1,046) of these showed a statistically significant rank-order correlation between the phylogenetic and geographic distances among their members, indicating a distance-decay relationship that is consistent with increasing dispersal limitation (Fig. 2B). Further, more diverse protein clusters were less likely to exhibit distance-decay relationships (Fig. 2C). Protein clusters with higher levels of diversity may be associated with a wider variety of viruses that are able to maintain their abundances across different habitat types and host availabilities, thus promoting their persistence across distances. Alternatively, the probability that a viral gene will overrun its own dispersal footprint, either by migrating around the planet or by doubling back on itself, may simply increase with time.

**Figure 2.**
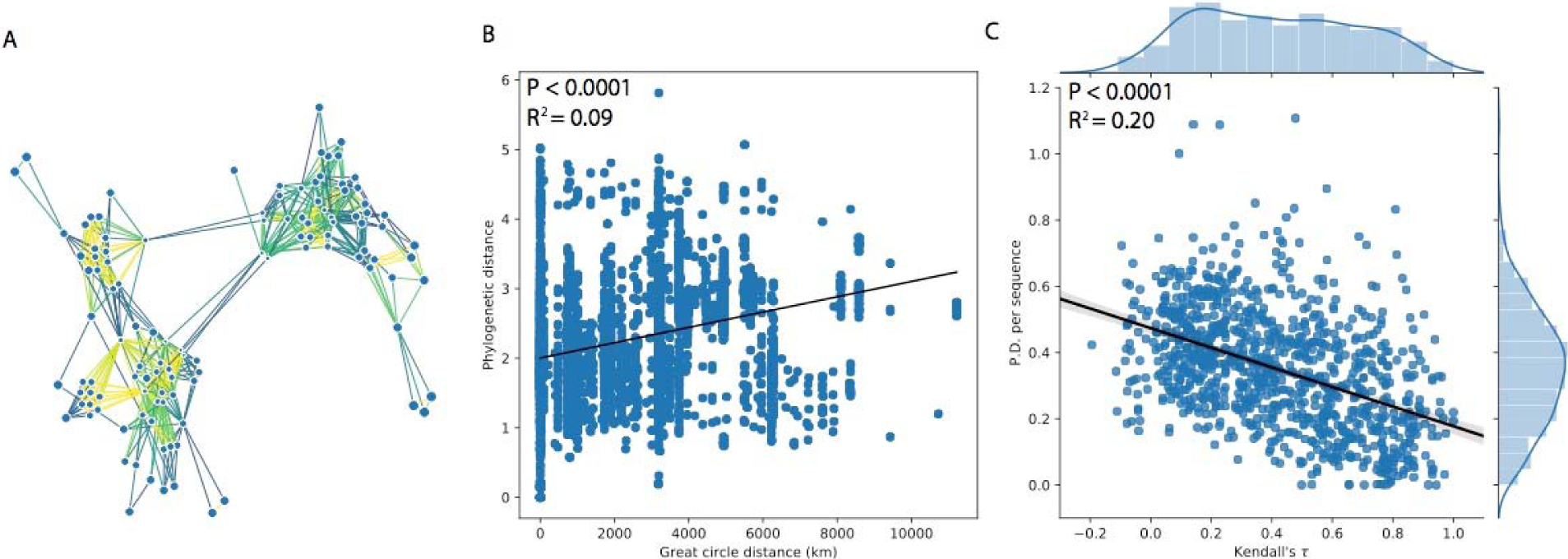
New viral protein clusters and distance-decay in sequence similarity. Example of distance-decay in protein clusters. Cluster 150 of 105,730 is shown for demonstration. (A) Network diagram of protein cluster 150 in which nodes represent predicted viral genes and edges denote sequence alignments. Edge colors correspond to bitscores, ranging from 48.3 (yellow) to 627.0 (blue). (B) Pairwise geographic distance of protein cluster 150 versus within-family phylogenetic distance. (C) Viral protein clusters whose phylogenies have a low phylogenetically weighted diversity (tree aspect ratio) tend to exhibit a more structured biogeography (β=-0.3, *R*^*2*^=0.2). P-values in (B) and (C) are derived from two-sided Wald tests.

## Microbial Hosts for Soil Viruses

We identified putative hosts of the soil viruses by assessing relationships between specific viral and microbial sequences. We first assigned hosts based on the similarity of CRISPR-spacer sequences to those found in microbial hosts deposited in the IMG/VR^32^ database. We also screened for similarities between viral contigs in our dataset and in viral isolates with known hosts. From these sequence-based approaches, we assigned microbial hosts to 208 viral contigs (0.8%). To extend this analysis, we also inferred virus-host relationships from co-occurrence networks containing both viral contigs and microbial OTUs (bacterial and archaeal) derived from 16 rRNA gene sequences in the metagenomes. When comparing these two approaches, we found that host assignment based on sequence homology substantially underestimated the number of viral hosts and yielded a distinct set of host organisms typically associated with pathogenesis, clinical applications, and/or marine environments (Fig. 3C). By contrast, correlations between viral contigs (Fig. 3A) and microbial sequences resulted in a more expansive range of putative host organisms associated with viruses. Only a small fraction of viral contigs had significant associations with specific microorganisms (440 of 19,094 contigs in the rarefied table; 2.3%), although many viral contigs (18,567) and microbial OTUs (12,512 of 21,895) were present at less than three locations and therefore excluded from our network (see methods). In the co-occurrence network, four viral contigs (0.9%) also had hosts identified by sequence homology, consistent with the proportion of sequence-identified hosts in the full dataset (0.8%). These contigs were predicted by sequence homology to target the same *Dickeya* species and were contained within a single cluster in the co-occurrence network, reinforcing similar host-relationships among these viruses (Fig. 3A, inset).

**Figure 3.**
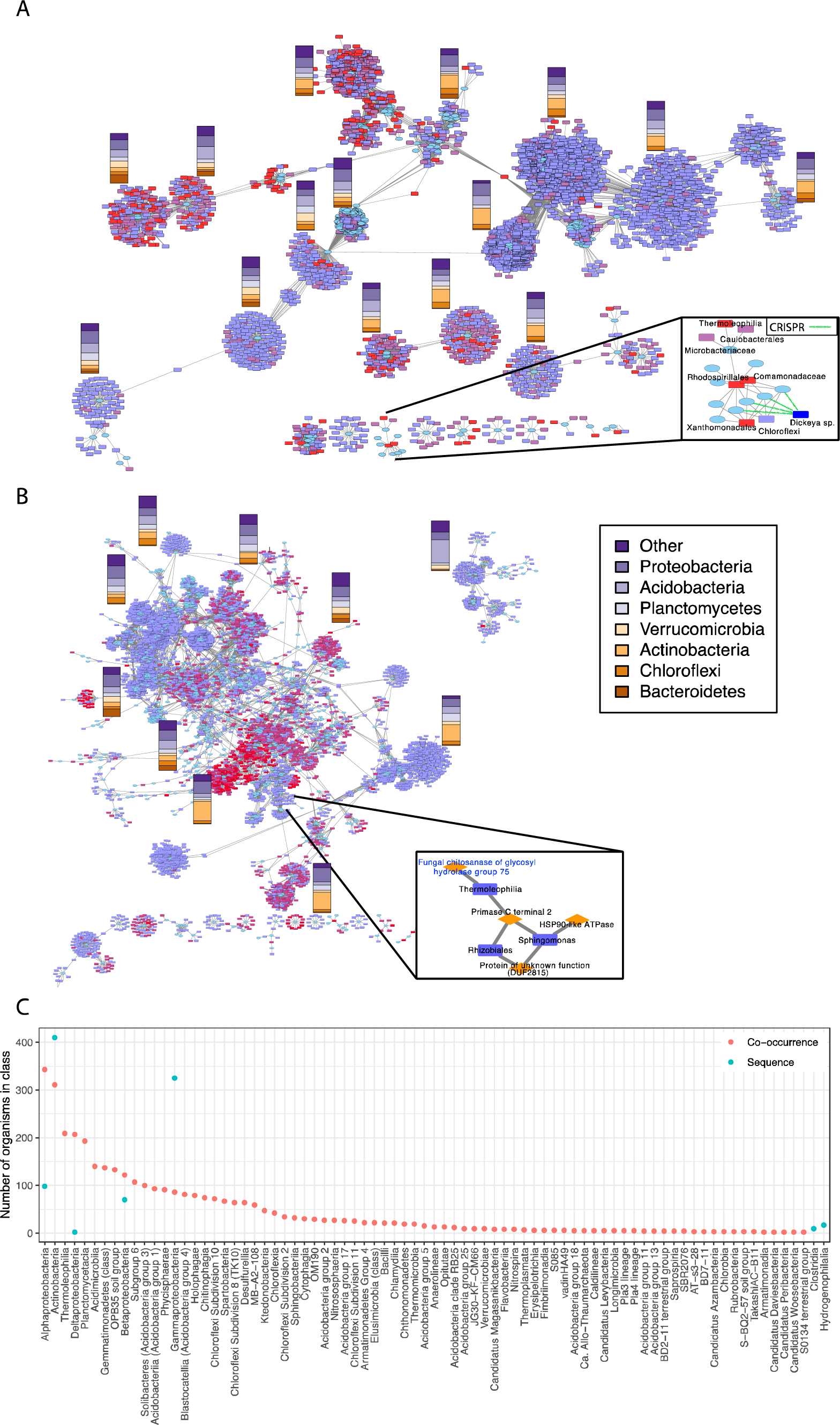
Soil virome host assignment through co-occurrence networks and sequence-based methods. Co-occurrence network with nodes representing viral or microbial OTUs and edges representing the co-occurrence relationships between them (*rho* > 0.6) is shown in Figure 2. Stacked bars indicate the relative percentages of microbial OTUs present in the adjacent clusters at the phylum level. Any grouping that had less than 10% representation in all clusters was rolled up to the “Other” category. Coloring of nodes indicates the relative abundance of that OTU in the cultivated prairie soil, with red indicating highest and blue indicating lowest abundance. (A) shows co-occurrences between viral sequences (ovals) and microbial OTUs (rectangles); (B) shows co-occurrences between viral pfams (ovals) and microbial OTUs (rectangles). The inset in (A) depicts a subnetwork with viral sequences (ovals) and microbial OTUs (rectangles), with the sequence-derived virus host relationships depicted with green dotted lines. The inset in (B) shows the cluster that contains a viral protein family containing a fungal chitosanase gene, which is discussed in detail below. Viral protein families are shown as diamonds in the inset and microbial OTUs are rectangles. (C) demonstrates the disparity in microbe-virus associations detected using co-occurrence networks versus sequence homology. Points denote the number of organisms in a given microbial class that were associated with viruses via each method.

More broadly, we also examined correlations between viral protein functional groups (pfams, Fig. 3B) and viral traits (e.g., tail, capsid, and membrane characteristics) with microbial OTUs. Using this trait-based approach, we obtained a much larger network of viral-host associations including correlations between 1,063 unique pfams and 5,665 unique microbial OTUs that grouped into 260 modules. Correlations between specific microorganisms and many viral traits were confined to a limited set of microorganisms per trait (79% of assigned modules contained 10 or fewer OTUs). However, the three largest modules each contained more than 250 interconnected pfams and microbial OTUs, and they were centered around viral traits for biosynthesis of vitamin B1 precursors (pfam13379), glycosyl transfer (pfam00953), and activation of the Hsp90 ATPase chaperone (pfam08327). The importance of these traits in co-occurrence network structure positions them as primary targets for deeper investigation viral-microbe relationships in soils, and more generally, we propose that trait-based approaches for studying the soil virome are beneficial for deciphering viral-host relationships in the highly diverse soil virome.

The network analysis also helped to identify potential broad host ranges in viruses, as most viral contigs showed significant positive correlations with phylogenetically-widespread microorganisms, consistent with recent work contrasting the historical paradigm of highly specific associations between viruses and microorganisms^3,33^. We identified 3,795 microbial OTUs as possible virus hosts via network analysis in contrast to 226 by sequence homology, over a ten-fold increase (Fig. 3A). Each viral contig was associated was 187 OTUs in the network on average, while individual viruses were assigned to a maximum of 19 possible hosts through sequence-based methods. Nevertheless, we detected some host specificity for some viruses because the microbial OTUs correlated with different viral contigs were distinct (mean Bray-Curtis dissimilarity = 0.89), as expected because specific viruses tend to infect certain groups of microorganisms^34^. The narrow set of hosts identified by sequence homology reveals a shortcoming of annotations that are relevant in soil settings. Improving sequence-based detection methods for detecting viral hosts in soil ecosystems is needed, as evidenced by the expansion of virus-microbe relationships with the network approach used here.

## Auxiliary Metabolic Genes (AMGs)

Possible AMGs were identified by removing known viral-associated protein families (as determined by Pfam) from our viral contigs. This yielded a large number (3,761) of unique protein families that were possible AMGs Screening of the AMG families according to their representation in different biomes revealed 302 cosmopolitan AMGs found in grassland, forest, sediment, arctic, and rhizosphere samples in comparison to 1,796 AMGs present in only one of these biomes. Cosmopolitan AMGs encoded functions such as glycosyl hydrolase/transferases (GH), peptidases, and cellulases (Supplementary Table 6).

Viral genes belonging to GH families were widespread in soils, as we found 43 GH families totaling 7,632 occurrences and three GH families in the top ten most abundant AMGs in our dataset (Supplementary Table 7), supporting previous studies indicating GHs as a key aspect of soil carbon cycles^12,16^. Among these, GH25 (lysozymes), GH108 (lysozymes), GH16 (transglycosylases active on plant and marine compounds), GH5 (cellulases, endomannanases, and related enzymes), and GH26 (endomannanases) were the most abundant (Supplementary Table 8). Some GH families are known to be active in viral lytic cycles (rev. in Davies et al.^17^), and the abundance of GH25 and GH108 in soil metagenomes further delineate this role. However, it is notable that three of the five most abundant GH families have possible functions in soil decomposition processes. When considering the broader suite of 43 GH families found in soil metagenomes, we posit that many of these genes play roles in soil decomposition that are beyond typical viral lysis functions. For instance, Emerson et al.^12^ previously suggested that viral-encoded endomannanases mediate permafrost C cycling and confirmed their functional abilities. In our soil dataset, putative endomannanase genes belonging to GH5 and GH26 alone occurred 532 times and were found in rhizosphere, arctic, and aquifer sediment biomes. Regardless, the distribution and sheer number of GHs as a whole generate a new understanding of virus-encoded C cycling genes as a ubiquitous feature of soil ecosystems and a reservoir of biogeochemical function.

We also investigated the impact of agriculture on soil viral AMGs. We predicted that cultivation would shift the composition of soil viromes due to shifts in soil properties such as pH, total nitrogen content, and fertilizer application, as previously demonstrated^5,6,13,35,36^. Metagenomes were screened from paired native soils and cultivated soils that were previously shown to contain a diverse array of microbial GHs known to degrade plant-derived polysaccharides^37^. We observed a 65% reduction AMG richness, including a lower number of GHs in cultivated relative to uncultivated soil metagenomes (Table 1). The disruption of this functional reservoir by land cultivation adds a new aspect to consider when investigating loss of soil function due to agricultural practices.

**Table 1.** Pfams associated with native vs. cultivated soils.

|  | **Native** | **Cultivated** |
| --- | --- | --- |
| Samples | 13 | 8 |
| Pfam occurrences | 4311 | 2847 |
| Pfam Richness | 1647 | 572 |
| Average pfam abundance | 2.617486 | 4.977273 |
| No. of pfams unique to land use | 1017 | 78 |
| Occurrences of unique pfams | 2524 | 355 |
| No. of glycoside hydrolase pfams | 8 | 4 |
| Occurrences of glycoside hydrolase pfams | 22 | 23 |
| No. of fungi-associated pfams | 3 | 0 |
| Occurrences of fungi-associated pfams | 8 | 0 |

## A New Paradigm: Cross-domain Transfer of Biogeochemical Function

We found genetic evidence for viral facilitation of multi-domain linkages in soils by uncovering bacteriophage sequences encoding a fungal chitinolytic gene; possibly signaling the transfer of a gene involved in fungal metabolism into bacterial hosts by viral infection. Specifically, we observed a fungal chitosanase (pfam07335) encoded by bacteriophages and verified the similarity of the bacteriophage chitosanase sequences to bacterial and fungal reference sequences previously deposited in databases (Fig. 4). Reference sequences clustered into distinct groups, and viral sequences from all soil metagenomes were interspersed with both domains of reference sequences. Chitosanases hydrolyze chitosan, a polymer of glucosamine residues that is an intermediate in chitin degradation, and are widely distributed in soil microorganisms that modulate C and nitrogen cycling (rev. in Somashekar and Joseph^38^). Chitin itself is a component of some fungal cell walls and insect exoskeletons that are common in terrestrial environments^39^. We identified 36 bacteriophage contigs in the full dataset that contained this chitosanase gene (Extended Data Figs. 2 and 3) and these contigs were primarily from vegetated biomes (Extended Data Tables 2 and 3). Also, when analyzing the entire IMG/VR virus database^40^, viral chitosanases were exclusively found in soil or freshwater viruses (66% vs. 33% of the cases, respectively). This finding highlights the potential importance of viral-encoded chitosanase functions in bacteria in ecosystems where chitin is abundant. Additionally, the limited distribution of viral chitosanase underlines the underexplored functional diversity associated with viruses in soils. The viral chitosanase genes that we identified may enable bacterial hosts to have easier access to nutrients in the heterogeneous soil environment by degrading free polysaccharides typically associated with fungal metabolism or by parasitizing live fungi^41^.

**Fig. 4.**
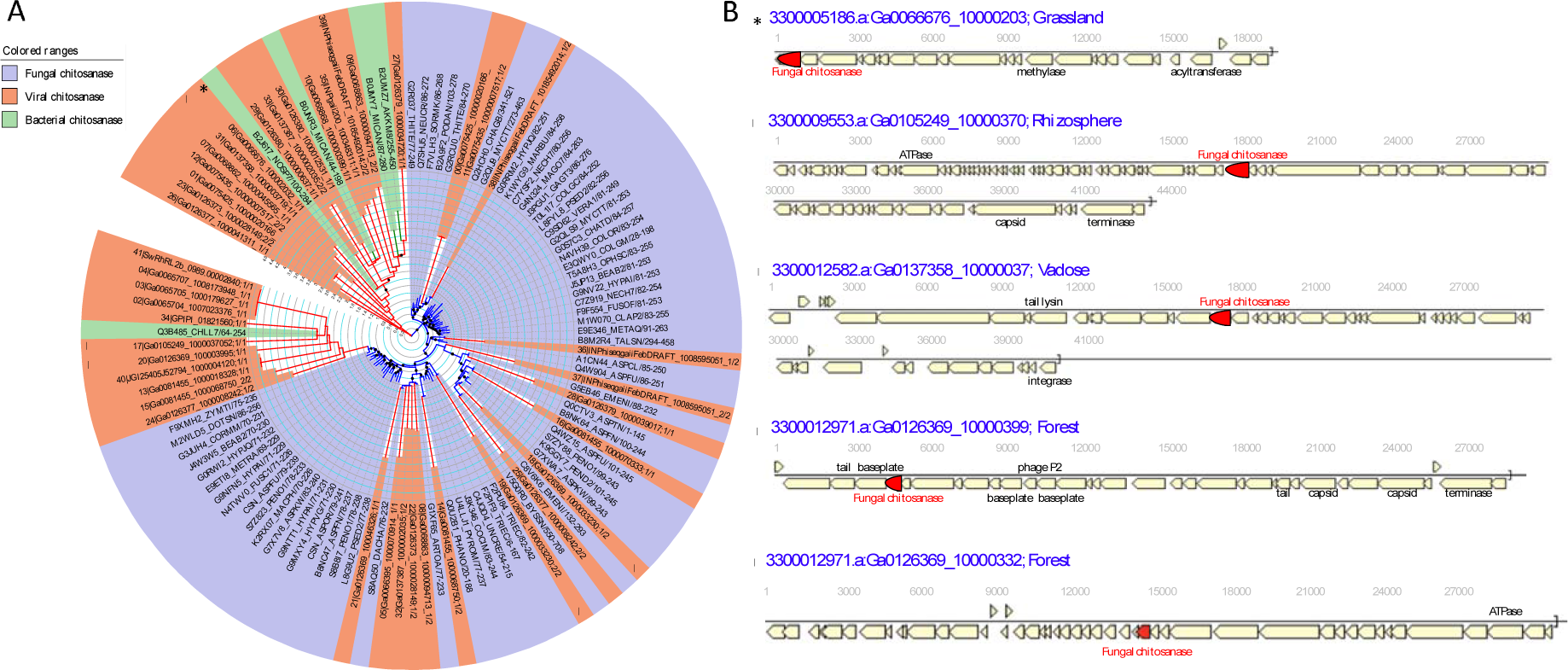
Alignment of soil viral chitosanase genes (pfam 07335) to existing chitosanase genes in bacterial and fungal databases. (A) Phylogenetic tree with chitosanase genes from the soil virome (orange) compared to genes found in bacterial (green) and fungal (purple) reference sequences. (B) Gene content of four representative bacteriophage contigs containing pfam07335. All contigs with pfam07335 are shown in Extended Data Fig. 2. Blue text: scaffold names of the virus contigs according to IMG/M and associated habitats. Predicted gene function is based on Clusters of Orthologous Genes (COG) categories. Chitosanase genes are shown in red. Symbols denote common sequences across panels (A) and (B)

Further support of a role for viruses in multi-domain metabolic exchange of soil C cycling genes include associations between viral sequences containing pfam07335 and microbial clades involved in decomposition within our co-occurrence network (Fig. 2B). Pfam07335 was the seed of a cluster containing three additional viral AMGs pfams encoding generic growth, replication, or unknown functions and three microbial OTUs belonging to *Actinobacteria, Rhizobiales*, and *Sphingomonas* (Extended Data Table 4). The co-occurrence of pfam07335 in viral sequences with microorganisms known to be major influencers of soil C cycling indicates that viruses may serve as both a reservoir encoding potential decomposition activities and a vector for the exchange of key soil functions across organisms. While unverified in the current work, such a relationship would be the first indication of cross-domain linkages in C cycling between bacteria, fungi, and viruses; and therefore, is a key area of future investigation.

## Conclusion

By mining over 24,000 viral metagenomic sequences from globally distributed terrestrial biomes, we reveal an incredibly diverse soil virome. We show that current computational methods for host assignment significantly underestimate possible viral-microbial interactions. Some viral proteins exhibit clear geographical distance-decay patterns similar to ecological patterns well-known in other soil-borne organisms. The soil virome contains a suite of cosmopolitan auxiliary metabolic genes (AMGs) encoding enzymes critical to decomposition. GHs, in particular, are highly abundant and co-occur with microorganisms mediating C cycling, thus lending genetic support to a biogeochemical role for soil viruses in exchanging decomposition metabolisms among key soil microorganisms. Finally, we show a fungal chitosanase found almost exclusively in soil bacteriophages in the uncultivated environment and vegetated environments more broadly. As a whole, our work exposes the soil virome as a reservoir of unexplored diversity that may be critical in the decomposition of soil organic matter and provides genetic evidence for viruses to aid in the transference of metabolic functions between bacterial and fungal domains.

## Materials and Methods

All statistical analyses were performed in *R* software version 3.3.1 using the packages ‘factoextra’^42^, ‘NbClust’^43^, ‘dplyr’^44^, ‘ggplot2’^45^, vegan^46^, and ‘gplots’^47^ unless otherwise noted.

### Viral sequence retrieval

All the viral sequences used in this work as well as their metadata (predicted host, viral grouping, taxonomy, and geographic location) were retrieved from the IMG/VR^32^ public data repository (https://img.jgi.doe.gov/vr/) version 1.0, a data management resource for visualization and analysis of globally identified metagenomic viral assembled sequences^3^ integrated with associated metadata within the IMG/M system^48^. All the viral sequences are over 5 kb. Both identification of the viral sequences and virus grouping were predicted using a computational approach fully described in Paez-Espino et al.^20^ in which an expanded and curated set of viral protein families was used as bait to identify viral sequences directly from metagenomic assemblies and highly related sequences (based on 90% identity over 75% of the alignment length on the shortest sequence) were clustered. For this work, we specifically mined the habitat type information of all the viral sequences and obtained 36,385 viral sequences under any of the categories: Terrestrial (soil), Terrestrial (other) and Host-associated (plants). We manually curated these datasets to remove obviously non-soil habitats (e.g., wastewater treatment reactors) to retain 24,334 viral sequences from 668 “soil-curated” samples. All viral sequences are annotated according to the DOE-JGI microbial genome annotation pipeline^49^. In addition, we have used pfams04451 and pfam16093 (hallmark protein families of major capsids of NCLDV viruses) and specific virophage major capsid protein models (Paez-Espino et al., in prep) to identify 1,676 giant virus sequences (42 > 5kb) and 538 virophage sequences (26 > 5kb), respectively, which are viral entities hardly identified with general discovery pipelines.

### Sequence-based host assignment for soil viruses

We used the host taxonomic information derived from IMG/VR version 1.0 where two computational approaches were used: (1) host assignment based on virus clusters that included isolate virus genomes with known hosts and (2) CRISPR-spacer sequence matches (only tolerating 1 SNP over the whole spacer length as cutoffs). To further complement the host assignment from IMG/VR version 1.0, we used a classification of the viral protein families (used in the virus identification pipeline) to determine the domain (Eukaryotic, Bacterial, or Archaeal) of the host predicted for 85.6% of the viral sequences described in Paez-Espino et al.^40^ (Supplementary Table 3). Briefly, the viral protein families were benchmarked against the viral RefSeq genomes and the viral genomes with predicted host from the prokaryotic virus of orthologous groups database^50^ obtaining a subset of them used as host-type marker genes.

### Pfam assignment

Structural and functional annotation of all sequences, including pfam assignment, is provided by the DOE Joint Genome Institute’s annotation pipelines^29^ where protein sequences are searched against Pfam database using HMMER 3.0^22^ using the gathering threshold (−-cut_ga) inside the pfam_scan.pl script^49^. That script also helps resolving potential overlaps between hits generating the final outcome. Sequences often contained multiple pfams, and we attributed each assigned pfam as occurring once per sequence.

### Read mapping of sequences against viral contigs

As described in Paez Espino et al.^20^ and applied in Paez Espino et al.^3^, we predicted the presence of any of the 24,334 soil viral sequences in low abundance across any of the 668 soil samples. We obtained all the assembled contigs and unassembled reads available from each of the soil samples and used the BLASTn program from the Blast+^51^ package to find hits (covering at least 10% of the virus length) to any of the predicted soil viral sequences with an e-value cutoff of 1e-5, a >=95% identity, and a > = 95% of the read/contig.

### Viral Abundance Quantification

Sequencing technology, depths, and sample numbers varied dramatically among studies. To account for these differences, we estimated viral loads by comparing the number of base pairs of sequence from raw reads that were attributed to viruses versus all base pairs in that metagenome. We evaluated the extent to which deeper sequencing would increase discoveries of viruses using linear regression of viral bp to bacterial bp. Fitting quadratic models to the relationship between bacterial and viral bp did not improve fit relative to linear models. Over- and under-representation of viral loads in each biome were using one-sample Student’s t-tests of residuals from linear models. We calculated average viral loads by multiplying the ratio of mean viral bp to mean microbial bp (unitless) by an estimate of average microbial bp per cell (bp/cell) and by the estimated average number of microbial cells per gram of soil (cell/g), yielding the number of viral bp per g of soil. We then divided by the average size bp per virus (bp/virus) in our dataset to derive the number of number of viruses per gram of soil. For comparison, we calculated viral loads in marine and human systems using the same procedure viral and microbial bp in raw metagenomic sequencing reads reported in Paez-Espino et al.^3^.

### Auxiliary Metabolic Gene Identification

Classifying AMGs is challenging due to the difficulty of defining genes that are external to viral replication and also allow viruses to manipulate host metabolism^16^. Recent work in soils has taken a targeted approach to exploring viral AMGs^37,52^. We started with viral-associated pfams as a basis for identifying AMGs. We filtered this list to remove known viral-specific pfams (Supplementary Table 9). We also searched for pfams whose annotations contained the following terms and removed all that were obviously viral: ‘phage’, ‘holin’, ‘capsid’, ‘tail’, ‘virus’, ‘viral’, ‘coat’, ‘lysis’, and ‘lytic’. Because GH genes are central in decomposition processes and common in soil bacteria^37,52^, we focused much of exploration on pfams containing genes in GH families. In total, we identified 3,761 unique AMGs present 653,536 times.

### Eco-Evolutionary Patterns

To assess distance-decay relationships in soil viromes, we first constructed viral protein clusters based on sequence similarity by using LAST to conduct an all-verses-all alignment of open reading frames from metagenomic viral contigs^53^. Alignment output was staged using ‘pandas’^54^. Then, a weighted undirected graph was populated using ‘networkx’^55^, with predicted viral genes represented as nodes, sequence alignments represented as edges and alignment bitscores represented as edge weights. Connected components were extracted and derived viral protein clusters as the sequences of within a single connected component.

The largest connected component contained a high proportion number of short alignments relative to other connected components. These short alignments appear to represent putative recombination events, convergence and coincidental alignments, so this large connected component was excluded from subsequent analysis. For each viral protein cluster, a multiple sequence alignment was performed using ‘Clustal Omega’^56^, and approximate maximum likelihood phylogenies were inferred using ‘fasttree’^57^. The geographic and phylogenetic distances were calculated using ‘SuchTree’^57^ and ‘Cartopy’^58^, and their correlation was estimated using the rank-order correlation coefficient and Kendall’s τ^59^. P-values were corrected for multiple testing using the Simes-Hochberg step-up procedure^60^.

### Bacterial and Archaeal 16S rRNA gene characterization

We used the high quality 16S rRNA identification and microbial prediction pipeline from the JGI^61^ (that uses a combination of Hidden Markov Models (HMMs) and sequence similarity-based approaches) based on complete/near complete gene sequence to obtain a grand total of 6,254 and 401,422 archaeal and bacterial 16 rRNA genes respectively across all the soil samples. Taxonomic information of this marker gene predicted lineages at different levels based on homology to the reference databases from domain to species as indicated.

We processed bacterial and archaeal sequences into operational taxonomic units (OTUs) as follows. Bacterial and archaeal sequences were assigned unique identifiers, followed by prefix dereplication using vsearch^62^. OTUs were clustered at 97% similarity, and reads were filtered out if they appeared only once or were flagged as chimeric by uchime de novo^62^. Reads were mapped to the filtered OTUs to construct an OTU table. BLAST+^51^ was used to align OTUs to Silva v128^63^ database and taxonomy is assigned using the CREST LCA method^64^.

### Co-Occurrence Networks

Because many microorganisms lack a CRISPR-Cas system^65^, we used co-occurrence networks to evaluate possible linkages between soil viruses and microorganisms (e.g., ^12,66-69^). We constructed two types of networks as described below – one correlating viral sequence abundance to microbial OTU abundance and one correlating pfams located on viral sequences to microbial OTU abundance. The purpose of the first network was to identify specific viral particles that were statistically co-located with specific microorganisms, while the second network revealed how viral traits (e.g., cap and tail physiology, modes of infection, AMGs, etc.) tended to be associated with certain microbial clades.

Sequencing depth and number of samples differed dramatically among samples (SI Table), so it was necessary to condense and rarefy data prior to network analysis. Each of three data types—viral sequences, viral pfams, and microbial OTUs—were processed independently of each other using the same workflow. To provide sufficient number of viral sequences to allow for rarefaction, we grouped all samples from the same location into a single data point by combining samples collected within 0.5° latitude and 0.5° longitude of each other. This increased sample sizes enough use abundance-based statistical approaches while allowing us to maintain habitat-specific differences in viromes and microbiomes. To generate robust rarefied datasets, we rarefied each data type 1000x and averaged counts across all tables to yield a final rarefied table. To choose the appropriate rarefaction level for each data type, we generated rarefaction curves for each data type and assessed reads per location at 10% intervals across the full read-per-location distribution. We also used histograms to visualize the number of reads per location and evaluate the number of locations that would be retained at each possible rarefaction level. Rarefaction levels for each data type were chosen to maximize both the number of sequences retained per location and the number locations retained. Our rarefaction process yielded 16 locations with 434 viral sequences, 25 locations with 715 viral pfams, and 14 locations with 1164 sequences microbial OTUs.

Co-occurrence networks were constructed using Spearman’s rank correlation coefficient as edges and both microbial OTUs and either viral sequences or pfams as nodes. Spearman correlations were calculated for all possible relationships, and only those with *rho* > 0.60 and an FDR-corrected p-value < 0.01 were included in networks. Correlations between two microbial OTUs or two viruses were not included in networks, such that only virus-microorganisms relationships are depicted. We also removed relationships between viral sequences or pfams with OTUs that were not co-located at least 3 times to prevent spurious correlations. Networks were visualized using Cytoscape version 3.6.1^70^. Modules were determined in Cytoscape using the FAG-EC^61^ algorithm in ClusterViz^71^ with selections set to ‘strong modules’ and a ‘ComplexSize’ threshold of 2. Differences in microbial OTUs across modules were evaluated with Bray-Curtis dissimilarity in the ‘vegan’ package in *R*.

### Viral chitosanase investigation

Sixty-nine well-curated seeds in the chitosanase pfam^21^ database (pfam07335) were divided into bacterial and fungal subsets based on the sequence dissimilarities and the taxonomy assignments obtained from NCBI taxonomy database^72^. We aligned the subset seeds and built bacterial and fungal HMMs separately with two iterations to obtain more robust models using HMMER v3.2.1^22^. The viral chitosanase domains were annotated by the HMM giving a higher bit score and retrieved from the alignments. A chitosanase reference tree was constructed using the 69 seed sequences via ‘FastTree’^73^. The viral chitosanase sequences were mapped to the reference tree based on the alignments to the HMM modeled positions without changing the tree topology using ‘pplacer’^74^. The fixed reference tree with the inserted branches of viral chitosanase sequences was visualized in iTOL v3^75^. Visualization of the gene content and gene location of the soil virus contigs containing viral chitosanases (using the gene neighborhood function from the IMG/M system) was used to verify their presence within the viral sequence (Extended Data Fig. 2).

